# Small molecule positive allosteric modulation of homomeric kainate receptors GluK1-3: Development of screening assays and insight into GluK3 structure

**DOI:** 10.1101/2023.11.02.565282

**Authors:** Yasmin Bay, Raminta Venskutonytė, Stine M. Frantsen, Thor S. Thorsen, Maria Musgaard, Karla Frydenvang, Pierre Francotte, Bernard Pirotte, Philip C. Biggin, Anders S. Kristensen, Thomas Boesen, Darryl S. Pickering, Michael Gajhede, Jette S. Kastrup

## Abstract

The kainate receptors GluK1-3 belong to the family of ionotropic glutamate receptors and are essential for fast excitatory neurotransmission in the brain and associated with neurological and psychiatric diseases. How these receptors can be modulated by small molecule agents is not well-understood, especially for GluK3. We show that the positive allosteric modulator BPAM344 can be used to establish robust calcium-sensitive fluorescence-based assays at GluK1-3 for testing agonists, antagonists, and positive allosteric modulators. The EC_50_ of BPAM344 for potentiating the response of 100 µM kainate was determined to 26.3 µM at GluK1, 75.4 µM at GluK2, and 639 µM at GluK3. In the presence of 150 µM BPAM344, domoate was found to be a potent agonist at GluK1 and GluK2 with EC_50_ of 0.77 µM and 1.33 µM, respectively. At GluK3, domoate acts as a very weak agonist or antagonist with IC_50_ of 14.5 µM, in the presence of 500 µM BPAM344 and 100 µM kainate. Using H523A mutated GluK3, we determined the first dimeric structure of the ligand-binding domain by X-ray crystallography, allowing location of BPAM344, zinc, sodium, and chloride ion binding sites at the dimer interface. Molecular dynamics simulations support the stability of the ion sites as well as the involvement of Asp761, Asp790, and Glu797 in binding of zinc ions. Using electron microscopy, we show that in the presence of glutamate and BPAM344, full-length GluK3 adopts a dimer-of-dimers arrangement. This study may contribute to unravelling the potential of kainate receptors as targets for treatment of brain diseases.

## Introduction

The fast synaptic excitatory effects of L-glutamate (Glu, Figure 1) in the mammalian central nervous system are mediated by signaling through the ionotropic glutamate receptors (iGluRs). As a result, these ligand-gated ion channels are essential for normal brain function but also play a role in a vast range of psychiatric and neurological disorders. The iGluRs have been classified into four families: the α-amino-3-hydroxy-5-methylisoxazole-4-propionate (AMPA) receptors, the kainate (KA, Figure 1) receptors, the N-methyl-D-aspartate (NMDA) receptors, and the Delta receptors [1]. Kainate receptors are expressed abundantly in the brain and have been related to several diseases, including epilepsy and depression [2]. In particular, the GluK3 subunit has been linked to major depression and schizophrenia on a genetic level [3, 4].

**Figure 1.**
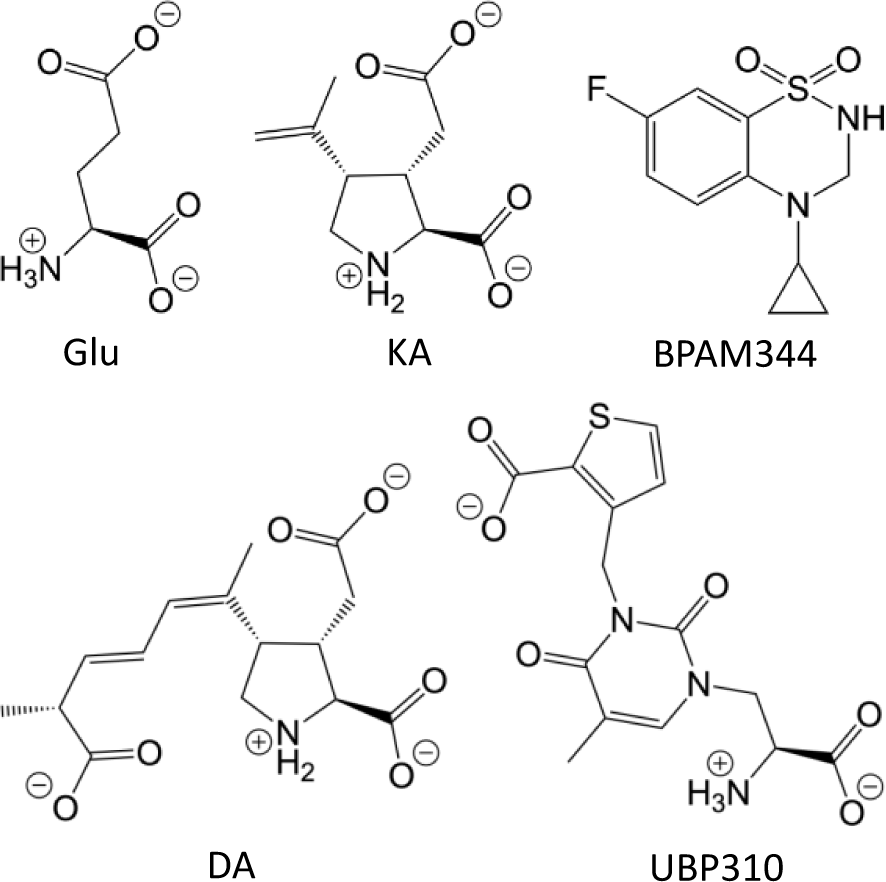
Chemical structures. The agonists L-glutamate (Glu), kainate (KA), and domoate (DA), the antagonist UBP310, and the positive allosteric modulator BPAM344, evaluated in the GluK1-3 calcium-sensitive fluorescence-based assays.

The iGluRs are composed of four subunits and share a common modular architecture [5]. Each subunit comprises a large extracellular domain layered into an amino-terminal domain (ATD) and a ligand-binding domain (LBD). The ion channel of the iGluRs is formed by all four subunits via a transmembrane domain (TMD) consisting of three membrane-spanning helices (M1, M3, and M4) and a re-entrant loop (M2). Lastly, each subunit has an intracellular carboxy-terminal domain (CTD) of various lengths. The LBDs form a dimer-of-dimers arrangement in the resting and activated state. Each LBD comprises two non-continuous polypeptide segments forming a clamshell structure with the glutamate binding site located in the cleft between the upper (D1) and lower (D2) lobes. Binding sites for positive allosteric modulators exist at the interface between two LBDs forming a dimer.

The kainate receptors can assemble from five subunits (GluK1-5) into either homomeric or heteromeric ion channels. GluK1-3 can form functional homomeric and heteromeric receptors, whereas GluK4-5 require co-assembly with GluK1-3 subunits to form functional heteromeric ion channels [6]. GluK3 is a puzzling receptor among the kainate receptors, as it has a very low sensitivity to glutamate with an EC_50_ in the millimolar range [7-9] compared to an EC_50_ in the micromolar range for GluK1 and GluK2 [10-12]. The low sensitivity to glutamate is due to fast desensitization of the GluK3 receptor at sub-saturating glutamate concentrations [7]; however, the functional implications of this low sensitivity are still unclear. In addition, GluK3 shows other intriguing pharmacological differences from the evolutionary-related GluK1 and GluK2 receptors, particularly its insensitivity towards domoate (DA, Figure 1) [9] and its sensitivity towards zinc ions [13]. This insensitivity towards domoate is further addressed in this study via the establishment of robust assays.

Anions and cations binding to allosteric sites in the LBD dimer interfaces have been shown to increase the glutamate response through stabilization of the LBD dimer, which increases the ability of the LBDs to translate agonist-binding to channel opening [14]. Specifically, it has been shown that activating kainate receptors by glutamate requires the presence of sodium and chloride ions [15]. In this context, a distinct property of the GluK3 receptor is that it is additionally modulated by extracellular zinc ions, which increase glutamate potency and delay desensitization by acting on Asp790 (numbering including signal peptide), which is unique to the GluK3 among the kainate receptor subunits [13]. In this study, we unravel potential binding sites for zinc, sodium, and chloride ions in GluK3.

The first structure of GluK3 in complex with glutamate was published in 2011 using a soluble construct of the ligand-binding domain of GluK3 (GluK3-LBD) [16], and later structures include complexes with glutamate and kainate [13, 17] as well as with synthetic agonists [18-22]. A characteristic of all GluK3-LBD structures is that the receptor crystallized as a monomer or a dimer that is not biologically relevant as the N-termini do not point in the same direction. In this study, we show the first structure of a biologically relevant dimer of GluK3.

Full-length structures determined by cryo-electron microscopy are now available of GluK1 [23], GluK2 [24-27], GluK3 [28, 29], and GluK2/K5 [30]. For GluK3, these include two structures of the full-length receptor determined in a closed state in the presence of the competitive antagonist UBP310 at 7.7 Å resolution [29] and UBP301 at 10.6 Å resolution [28]. Both structures appear to represent GluK3 in a partially “recovered” state with two LBDs in a dimeric arrangement, whereas the location of the other two LBDs resembled the location in the desensitized state. Additionally, two structures of GluK3 in the desensitized state have been determined in the presence of the agonist (2*S*, 4*R*)-4-methylglutamate (SYM) at 7.4 Å resolution [29] and kainate at 9.6 Å resolution [28] and show domain organization similar to that of GluK2. Recently, structures of GluK2 with BPAM344 in the closed state showed that two BPAM344 molecules bind per ligand-binding domain dimer interface [27], as previously reported using a soluble construct of the ligand-binding domain of GluK1 [31]. In this study, we show that it is possible to capture GluK3 in a dimer-of-dimers arrangement, resembling the active state.

While numerous positive allosteric modulators of AMPA receptors are available [32], only recently small-molecule positive allosteric modulators of GluK1-3 have been reported [31, 33] and their binding site determined in GluK1-LBD [31]. Of particular interest, the 1,2,4-benzothiadiazine 1,1-dioxide modulator BPAM344 (Figure 1) was shown to dramatically potentiate glutamate-evoked currents of GluK3_a_ as well as GluK2_a_, GluK1_b_, and GluA1_i_ in a concentration-dependent manner. In contrast, the plant lectin concanavalin A (ConA), which has been employed in kainate receptor research to reduce desensitization of GluK1 and GluK2 receptors, has a very limited effect on GluK3 receptors [9]. In this study, we show how BPAM344 binds to and modulate the structure of GluK3.

Our first aim was to establish robust assays at GluK1-3 that would allow testing of agonists, antagonists, and positive allosteric modulators. For this purpose, we established calcium-sensitive fluorescence-based assays for the homomeric GluK1-3 kainate receptors, using BPAM344 as a reference compound to diminish desensitization of the receptors. The results show that the assays allow for the screening of novel tool compounds that can aid in the development of kainate receptor selective agonists, antagonists, and positive allosteric modulators. The second aim was to investigate how BPAM344 and ions, such as zinc, sodium, and chloride ions, bind to the less well-understood kainate receptor GluK3 using X-ray crystallography and molecular dynamics simulations on the ligand-binding domain of GluK3 and electron microscopy on full-length GluK3. We report an X-ray structure at 2.7 Å resolution of the ligand-binding domain of GluK3 with kainate. The receptor was crystallized as a biologically relevant dimer by introducing the H523A mutation (GluK3-H523A-LBD). Furthermore, we determined the structure of GluK3-H523A-LBD with kainate plus BPAM344 at 2.9 Å resolution. These structures allowed the identification of BPAM344 binding sites at the LBD dimer interface as well as binding sites for cations and anions. The third aim was to address if it would be possible to capture GluK3 in a dimer-of-dimers arrangements in the presence of a positive allosteric modulator. Using electron microscopy, we show that in the presence of glutamate and BPAM344, full-length GluK3 can adopt a dimer-of-dimers arrangement as previously reported for the AMPA receptor GluA2 [34].

## Results

### Establishment of calcium-sensitive fluorescence-based assays

Stable GT-HEK293 cell lines expressing GluK1, GluK2, or GluK3 were produced. The cell lines were validated using homologous competition binding with [^3^H]-NF608 (16.3 Ci/mmol) as radiolabel for GluK1 and [^3^H]-kainate (67.5 Ci/mmol) as radiolabel for GluK2 and GluK3 (Figure S1). The radioligand affinities were in the nanomolar range: K_d_ = 33 nM, 23 nM, and 20 nM for GluK1, GluK2, and GluK3, respectively. These values are within the same range as previously reported [35, 36].

For initial validation of the stable cell lines, the ability of glutamate and two well-known kainate receptor agonists, kainate and domoate, to induce Ca^2+^ influx in the presence or absence of 300 µM BPAM344 was tested. BPAM344 significantly potentiated the responses evoked by glutamate, kainate, and domoate at GluK1 and GluK2 (Figure 2(A-B)). At GluK3 expressing cells, BPAM344 potentiated glutamate and, to an even greater extent, kainate-induced responses (Figure 2(C)). In contrast, BPAM344 did not potentiate GluK3 responses to domoate (*P*-value 0.165 for 1 µM and 0.484 for 10 µM domoate).

**Figure 2.**
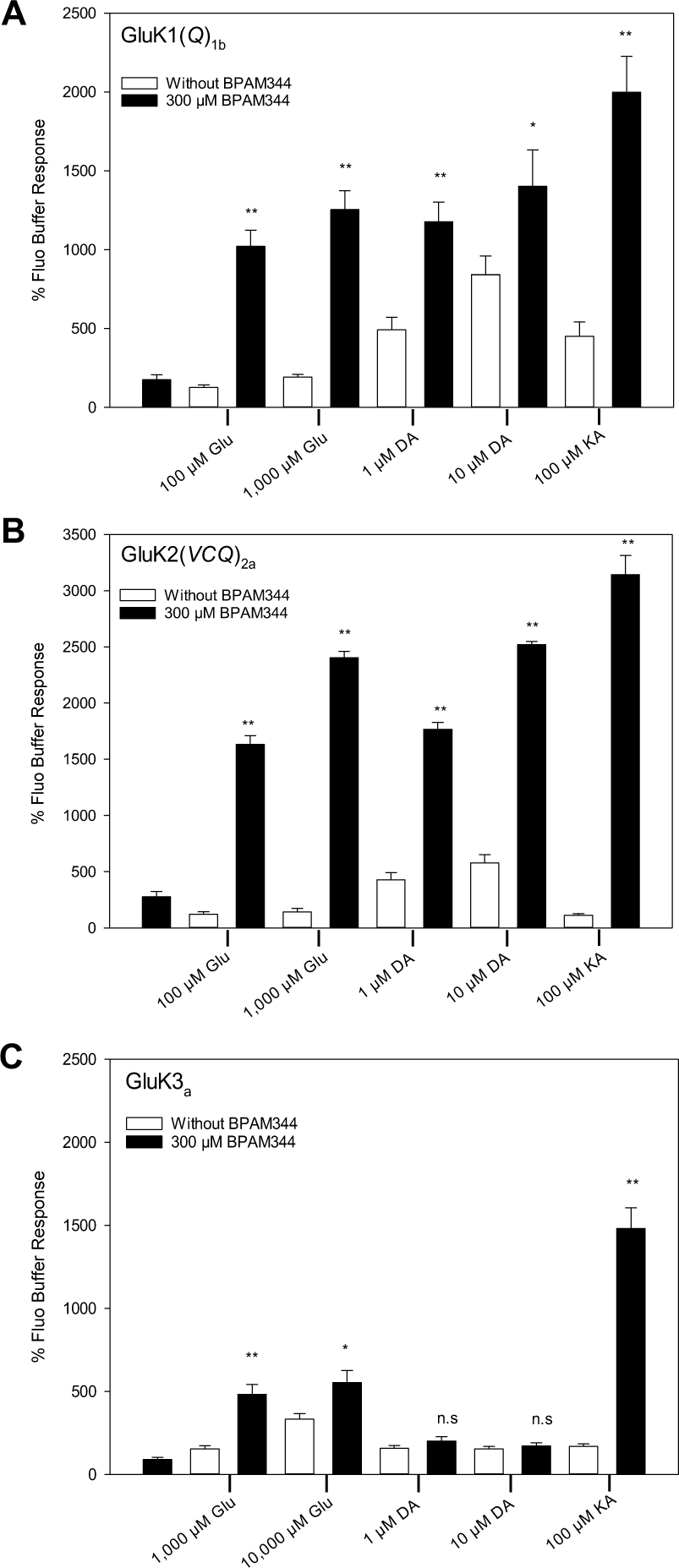
Evaluation of the BPAM344 modulatory effect in the presence of three agonists at the stable cell lines expressing the homomeric kainate receptors GluK1-3. BPAM344 response with the agonists glutamate (Glu), kainate (KA), and domoate (DA) at GT-HEK293 cells stably expressing (A) GluK1(*Q*)_1b_, (B) GluK2(*VCQ*)_2a_, and (C) GluK3_a_. Data are normalized to the response of Fluo buffer and shown as mean ± SEM. Student’s t-test was used for assessment of responses in the presence or absence of BPAM344: ** (*P* < 0.001), * (*P* = 0.009-0.022), and n.s, not significant (*P* = 0.165-0.484). *P* values equal to or between 0.001-0.022 were obtained at GluK1, GluK2, and GluK3 for Glu, DA, and KA, except for DA at GluK3.

### Potency of glutamate, kainate, and domoate in the calcium-sensitive fluorescence-based assays

The potency (EC_50_) of the agonists (glutamate, kainate, and domoate) at each receptor was determined in the presence of 150 or 500 µM BPAM344. At the cell line expressing GluK1, nano-to micromolar potencies were observed with the rank order of potency being domoate>kainate>glutamate (Figure 3(A), Table 1), which is in accordance with the rank order previously reported using the potentiator ConA to block desensitization [10-12]. The same rank order was observed in GluK2 expressing cells (Figure 3(B), Table 1), and in agreement with the rank order previously reported in the presence of ConA [10-12]. At the GluK3 expressing cell line kainate was likewise more potent than glutamate (Figure 3(C), Table 1). An upper plateau of the concentration-response curve of glutamate was not reached even at 100 mM; thus, the determined EC_50_ value of 24 mM approximates the true potency. The EC_50_ of glutamate at GluK3 is comparable to the potency of 5.9-12.4 mM determined by electrophysiology [7-9]. However, it is worth noting that these values were also determined from incomplete curves using only up to 30 mM glutamate, indicating that the actual EC_50_ value could be higher.

**Figure 3.**
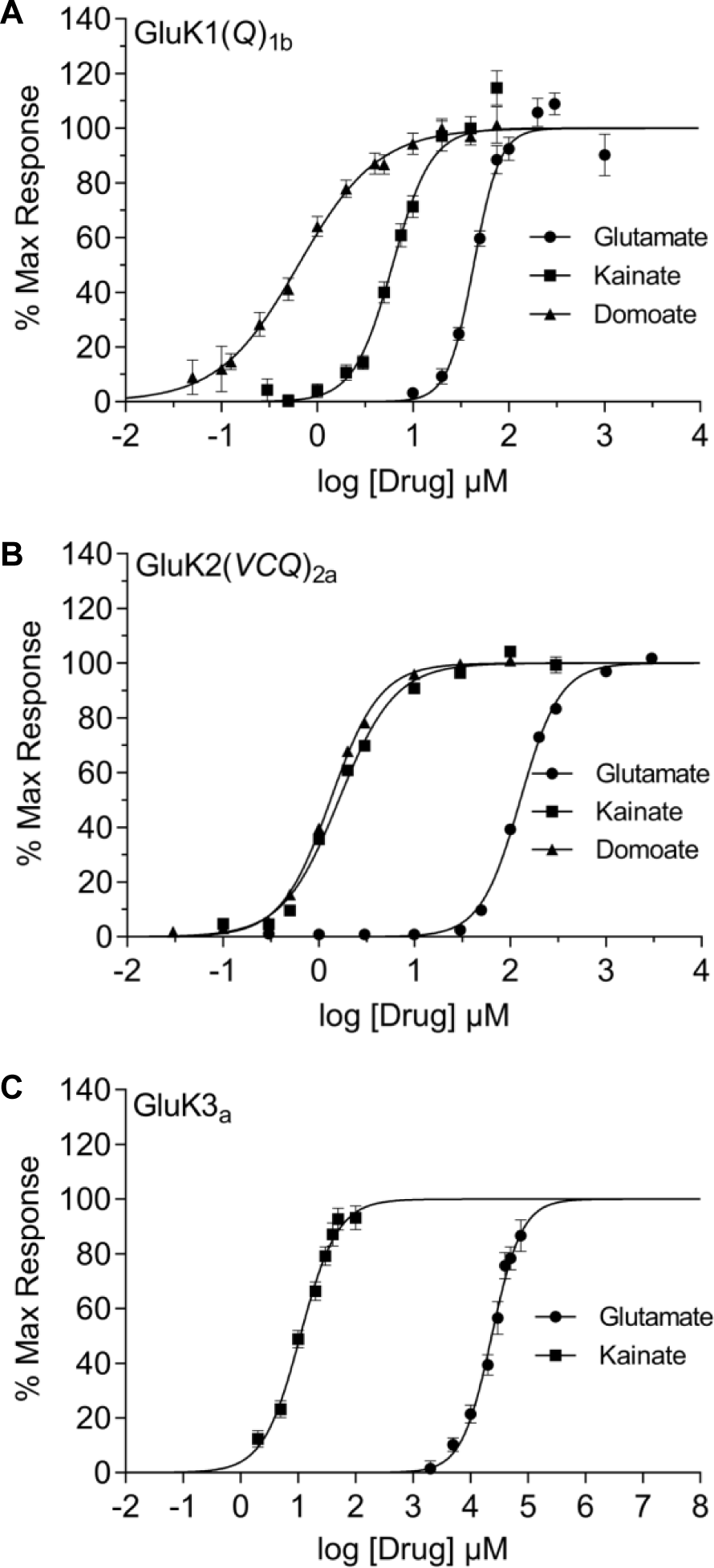
Pharmacological profile at the homomeric kainate receptors GluK1-3 in response to glutamate, kainate, and domoate. Concentration-response curves of the agonists glutamate, kainate, and domoate at GT-HEK293 cells expressing (A) GluK1(*Q*)_1b_, (B) GluK2(*VCQ*)_2a_ and (C) GluK3_a_. The concentration of BPAM344 was 150 or 500 µM (see Table 1). The data points for each curve are expressed as the mean ± SEM of octuplicate wells (n = 8). One concentration-response curve represents pooled data from 3-4 independent experiments by normalizing the raw relative fluorescence unit (RFU) values to the maximum response and constraining the bottom of the curve to 0% and the top of the curve to 100%. EC_50_ values are given in Table 1.

**Table 1.** Calcium-sensitive fluorescence-based assays. Functional evaluation of agonists, antagonists and BPAM344 at the stable cell lines expressing GluK1(*Q)_1b_,* GluK2(*VCQ*)_2a_, and GluK3_a_.

| Compounds | GluK1(Q)<sub>1b</sub> | GluK2(VCQ)<sub>2a</sub> | GluK3<sub>a</sub> |
| --- | --- | --- | --- |
|  | Potency (μM)* | Potency (μM)* | Potency (μM)* |
| **Glutamate** | 43 ± 8<sup>†</sup> | 129 ± 2<sup>†</sup> | 23900 ± 790<sup>†</sup> |
| **Kainate** | 6.6 ± 1.0<sup>†</sup> | 1.66 ± 0.07<sup>†</sup> | 11.4 ± 0.5<sup>†</sup> |
| **Domoate** | 0.77 ± 0.13<sup>†</sup> | 1.33 ± 0.03<sup>†</sup> | 14.5 ± 1.0<sup>‡</sup> |
| **UBP310** | 0.0200 ± 0.0041<sup>§</sup> | n.d. | n.d. |
| **BPAM344** | 26.3 ± 5.1<sup>¶</sup> | 75.4 ± 4.7<sup>¶</sup> | 639 ± 29<sup>¶</sup> |

\* Mean EC<sub>50</sub> or IC<sub>50</sub> ± SEM. The data are obtained from 3-4 independent experiments performed in octuplicate for each compound.
<sup>†</sup> EC<sub>50</sub> value ± SEM was determined in the presence of 150 μM BPAM344 for glutamate, kainate, and domoate at the GluK1 and GluK2 cell lines, and glutamate and kainate at the GluK3 cell line.
<sup>‡</sup> IC<sub>50</sub> value ± SEM was determined for domoate at the GluK3 cell line in the presence of 500 μM BPAM344 and 100 μM kainate.
<sup>§</sup> IC<sub>50</sub> value ± SEM was determined for UBP310 at the GluK1 cell line in the presence of 500 μM BPAM344 and 100 μM kainate.
<sup>||</sup> n.d. = not determined.
<sup>¶</sup> EC<sub>50</sub> value ± SEM of BPAM344 was determined in the presence of 100 μM kainate at all three cell lines.

### Potency of antagonists in the calcium-sensitive fluorescence-based assays

To investigate if the assay set up could also be used for testing the potency of antagonists, an inhibition curve for the selective GluK1 antagonist UBP310 was determined using 100 µM kainate and 500 µM BPAM344. The determined IC_50_ value of 0.02 µM (Figure 4(A), Table 1) agrees with the previously reported IC_50_ value of 0.01 µM [37]. Thus, the calcium-sensitive fluorescence-based assays can be used to screen for new antagonists at kainate receptors.

**Figure 4.**
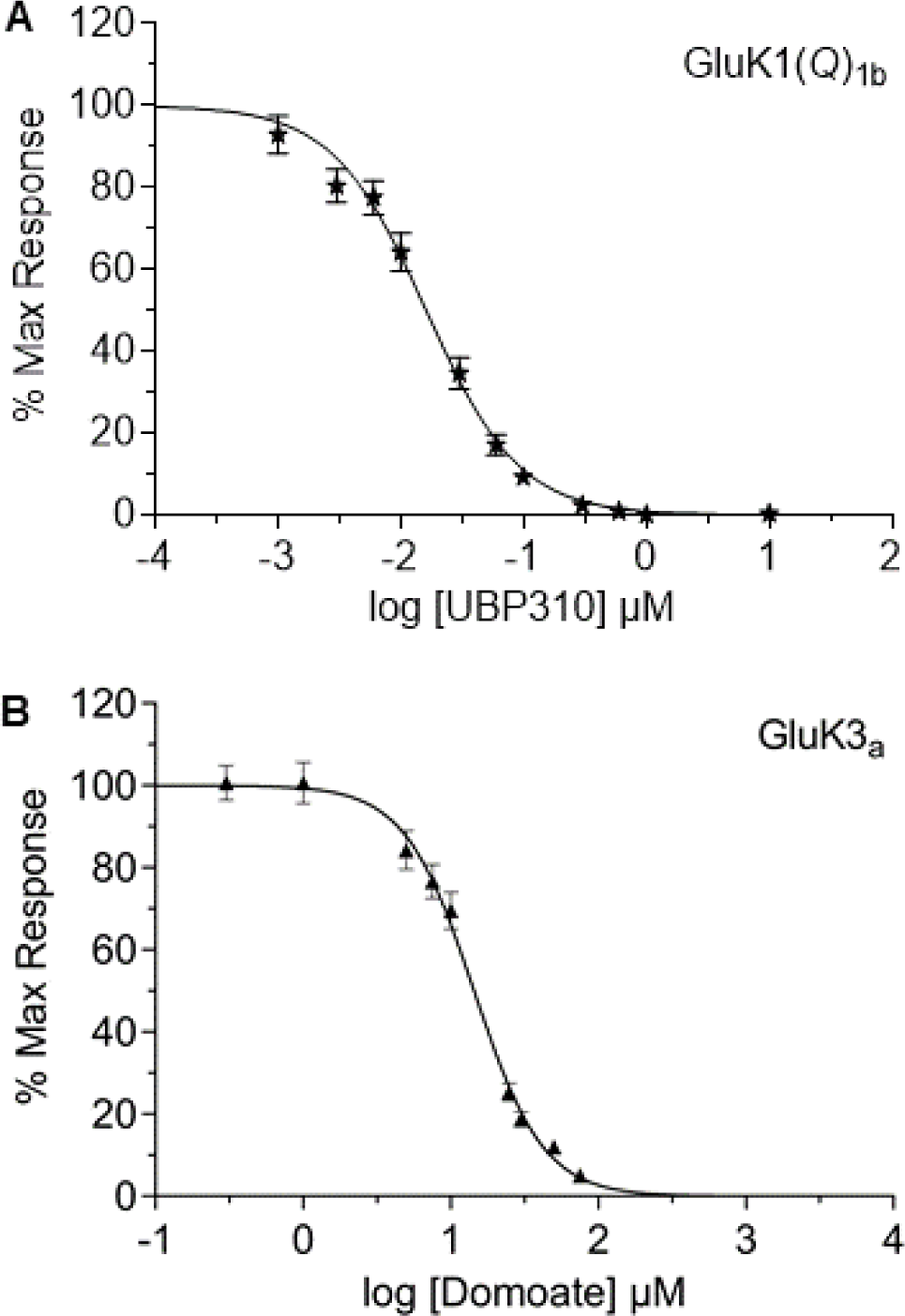
Inhibition curves at the homomeric kainate receptors GluK1 and GluK3 in response to the selective GluK1 antagonist UBP310 and domoate, respectively. Concentration-response curves of (A) UBP310 at GT-HEK293 cells expressing GluK1(*Q*)_1b_ and (B) domoate at GT-HEK293 cells expressing GluK3_a_, both in the presence of 500 µM BPAM344 and 100 µM kainate. The data points for each curve are expressed as the mean ± SEM of octuplicate wells (n = 8). One concentration-response curve represents pooled data from 3-4 independent experiments by normalizing the raw RFU values to the maximum response and constraining the bottom of the curve to 0% and the top of the curve to 100%.

Due to lack of response to domoate at GluK3, domoate was instead investigated as an antagonist, yielding an IC_50_ of 14.5 µM (Figure 4(B), Table 1). These results indicate that domoate acts as a very weak partial agonist at GluK3 receptors.

### Potency of BPAM344 in the calcium-sensitive fluorescence-based assays

The potency of BPAM344 at each kainate receptor was determined in the presence of 100 µM kainate (Figure 5, Table 1). The greatest potency of BPAM344 was observed at GluK1 (EC_50_ of 26.3 µM), whereas an EC_50_ of 75.4 µM was observed at GluK2. The EC_50_ of BPAM344 at GluK2 is very well in agreement with the 79 µM potency estimated by Larsen *et al*. in electrophysiological experiments [31]. BPAM344 has the weakest potency at GluK3 (EC_50_ of 639 µM).

**Figure 5.**
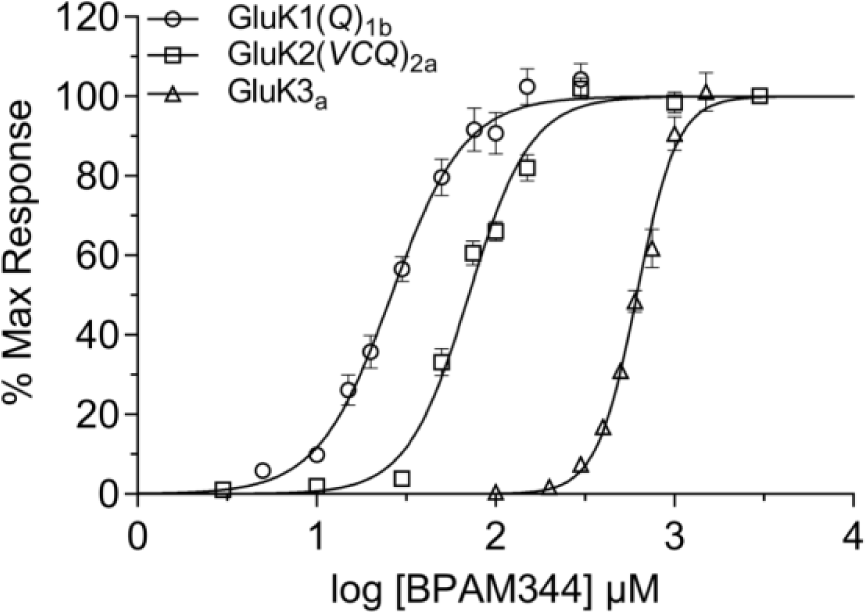
Pharmacological characteristics of the GluK1-3 stable cell lines, showing a distinct modulatory effect of BPAM344 at the different subtypes. Concentration-response curves for BPAM344 at GT-HEK293 cells expressing either GluK1(*Q*)_1b_, GluK2(*VCQ*)_2a_, or GluK3_a_ in the presence of 100 µM kainate. For graphical representation of each concentration-response curve, responses were pooled from 3-4 experiments and the average raw RFU at a given drug concentration was normalized to calculate the maximum response. Curves were constrained to bottom (0%) and top (100%).

### Biologically relevant dimeric structure of GluK3-H523A-LBD

To facilitate crystallization of the biological dimer, we mutated His523, located at the dimer interface, into an alanine to prevent this residue from interfering with crystal packing contacts as previously observed [13], and then expressed the mutant protein in *E. coli*. The X-ray crystal structure was solved at 2.7 Å resolution with two dimers of the GluK3-H523A-LBD in the asymmetric unit of the crystal (Table 2). These GluK3-H523A-LBD dimers (Figure 6(A)) resemble the biologically relevant iGluR dimers previously seen in other LBD structures of AMPA and kainate receptors [38-41]. Ala523 is located at the dimer interface, facing Ile781of the adjacent subunit, which together with a neighboring Leu784 creates a hydrophobic pocket within the dimer interface (Figure 6(B)).

**Figure 6.**
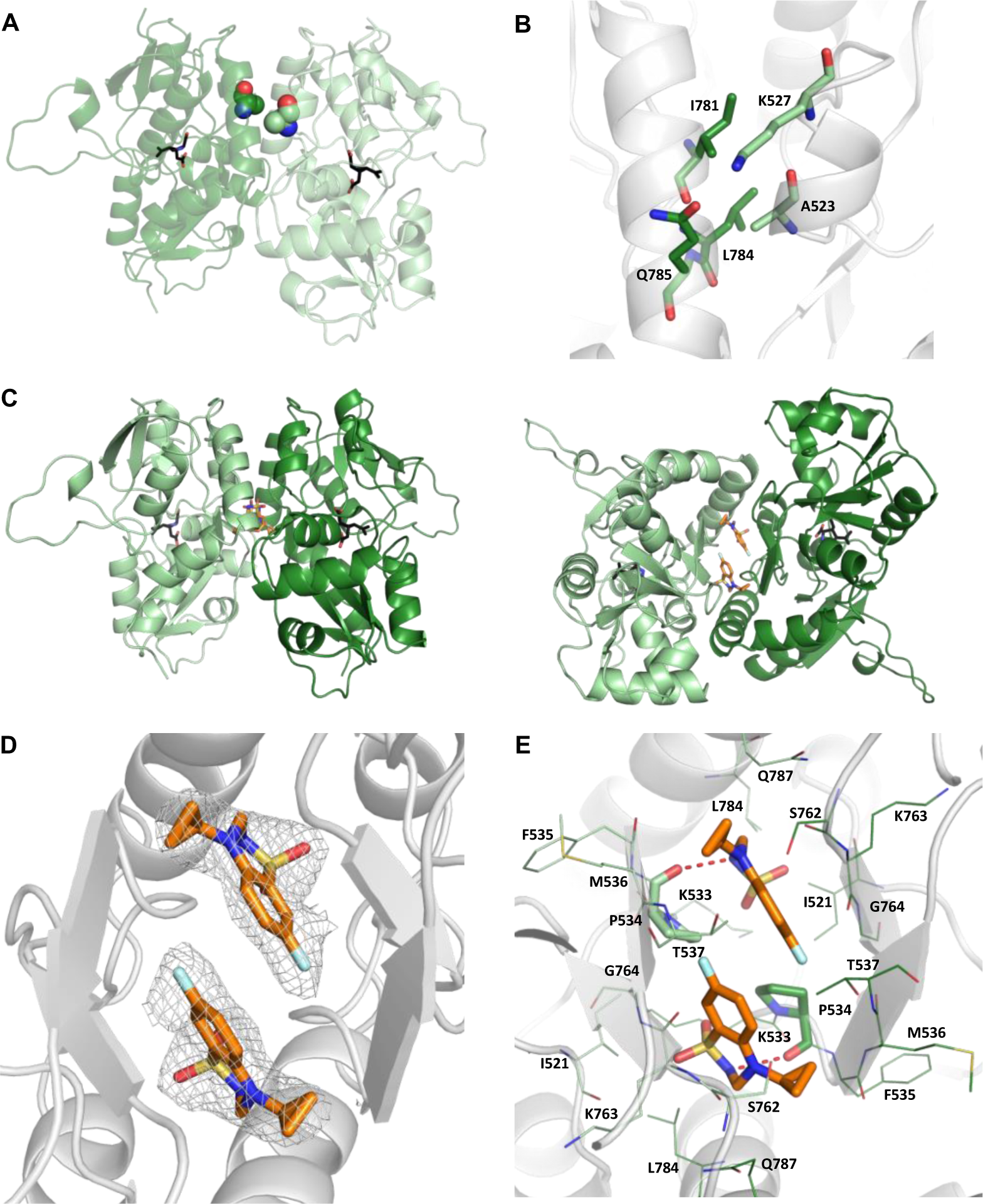
GluK3-H523A-LBD dimer with kainate and BPAM344. (A) Cartoon representation of the GluK3-H523A-LBD dimer with kainate. Chains A and B are shown in light and dark green, respectively. Kainate is shown in sticks representation with carbon atoms in black and Ala523 at the dimer interface as spheres representation. (B) Zoom on the H523A site at the dimer interface with surrounding residues shown in sticks representation. Oxygen atoms are red and nitrogen atoms blue. (C) Cartoon representation of the GluK3-H523A-LBD dimer with kainate and BPAM344. Two different views are shown, illustrating that two molecules of BPAM344 bind at the dimer interface. Left: side view with lobes D1 at the top, Right: bottom view. Chains A and D are displayed in light and dark green, respectively. BPAM344 is shown with carbon atoms in orange sticks representation and kainate as black sticks. (D) Simple 2Fo-Fc electron density map contoured at 1 sigma and carved 2.0 Å around BPAM344 in light grey. (E) Zoom on the positive allosteric modulator binding sites. Residues within 4 Å of the two BPAM344 molecules are shown in lines representation, except for Pro534 that is shown as sticks. The hydrogen bonds from BPAM344 to Pro534 are shown as red dashed lines. Oxygen atoms are red, nitrogen atoms blue, sulfur atoms yellow, and fluorine atoms light blue.

**Table 2.** X-ray data collection and refinement statistics for GluK3-H523A-LBD.

| Data collection | Kainate | Kainate and BPAM344 |
| --- | --- | --- |
| Space group | $P2_1$ | $P2_1$ |
| Cell dimensions a, b, c (Å), $\beta$ (°) | 69.41, 101.60, 83.41, 111.99 | 83.15, 100.56, 132.83, 103.93 |
| Resolution (Å)* | 42.46-2.70 (2.83-2.70) | 46.84-2.90 (3.06-2.90) |
| Completeness (%)* | 99.7 (99.5) | 99.9 (99.7) |
| $I/\sigma I$* | 4.3 (0.9) | 6.3 (1.8) |
| Redundancy<sup>a</sup> | 4.3 (4.3) | 4.1 (4.1) |
| CC1/2* | 0.98 (0.72) | 0.99 (0.85) |
| $R_{\text{merge}}$ (%)<sup>a</sup> | 11.5 (71.0) | 11.6 (43.3) |
| **Refinement** |  |  |
| No. reflections (work / R-free set) | 29483/1468 | 47131/2378 |
| $R_{\text{work}} / R_{\text{free}}$ (%) | 0.2036/0.2377 | 0.2068/0.2345 |
| No. residues |  |  |
| Protein A/B/C/D/E/F/G/H | 252/253/242/247/-/-/-/- | 252/247/252/253/247/248/253/248 |
| Kainate/BPAM344 | 4/- | 8/8 |
| Zn/Cl/SO<sub>4</sub>/glycerol/acetate/PEG/water | 13/7/10/5/6/1/97 | 20/22/8/13/3/-/115 |
| $B$-factors (Å<sup>2</sup>) | | |
| Protein | 54.1/56.3/68.2/58.6/-/-/-/- | 39.5/32.8/38.7/33.6/33.0/37.9/46.5/34.2 |
| Kainate/BPAM344 | 43.0/- | 29.0/23.2 |
| Zn/Cl/SO<sub>4</sub>/glycerol/acetate/PEG/water | 72.6/59.1/68.9/55.4/55.4/59.9/44.2 | 41.9/42.0/42.3/31.5/41.1/-/30.8 |
| R.m.s. deviations |  |  |
| Bond lengths (Å) | 0.002 | 0.002 |
| Bond angles (°) | 0.474 | 0.491 |
| Ramachandran outliers (%) | 0.0 | 0.0 |

\* Values in parentheses are for highest-resolution shell.

### BPAM344 binding sites at the dimer interface

BPAM344 has previously been crystallized in the ligand-binding domain of GluK1 (GluK1-LBD), revealing two binding sites for BPAM344 at the dimer interface [31]. To localize the BPAM344 binding site(s) in GluK3, we crystallized GluK3-H523A-LBD with kainate and BPAM344. The structure was determined at 2.9 Å resolution and contained four dimers in the asymmetric unit. Kainate binds at the orthosteric binding site in all eight molecules, whereas two molecules of BPAM344 were found at each dimer interface in all four GluK3-H523A-LBD dimers (Figure 6(C-D)), in a site located around a pseudo two-fold symmetry axis as previously seen in GluA2-LBD [38] and GluK1-LBD [31].

The binding mode of BPAM344, as well as binding site residues, is very similar to that in GluK1-LBD [31]. One direct hydrogen bond is established from the sulfonamide N-H proton of BPAM344 to the backbone O atom of Pro534 (Figure 6(E)). Furthermore, several van der Waals interactions between BPAM344 and surrounding residues (within 4 Å) from both subunits are observed to Lys533, Pro534, Phe535, Met536, Thr537, Leu784, and Gln787 from one subunit and Ile521, Pro534, Met536, Thr537, Ser762, Lys763, and Gly764 from the other subunit. All these residues are conserved between GluK1 and GluK3.

### Ion binding sites at the dimer interface

A chloride ion was located at the dimer interface, interacting with Lys533 in both subunits. An ion at this site was observed in both the structure with and without BPAM344 (Figure 7(A-C)). The residues comprising the sodium binding site are also the same in GluK1-3 (Figure 7(D)). However, probably due to a medium resolution of the structures, it was not possible to unravel if the electron density present at this site corresponds to a sodium ion or a water molecule. When performing molecular dynamics (MD) simulations on the GluK3-H523A-LBD structure without sodium or chloride ions present at these sites, sodium ions rebind to at least one of the two sites (Figure S2), supporting that the sodium sites are truly conserved. Furthermore, when initiating simulations from a structure with two sodium ions and one chloride ion bound, all three ions bind stably throughout the simulations (Figure S2).

**Figure 7.**
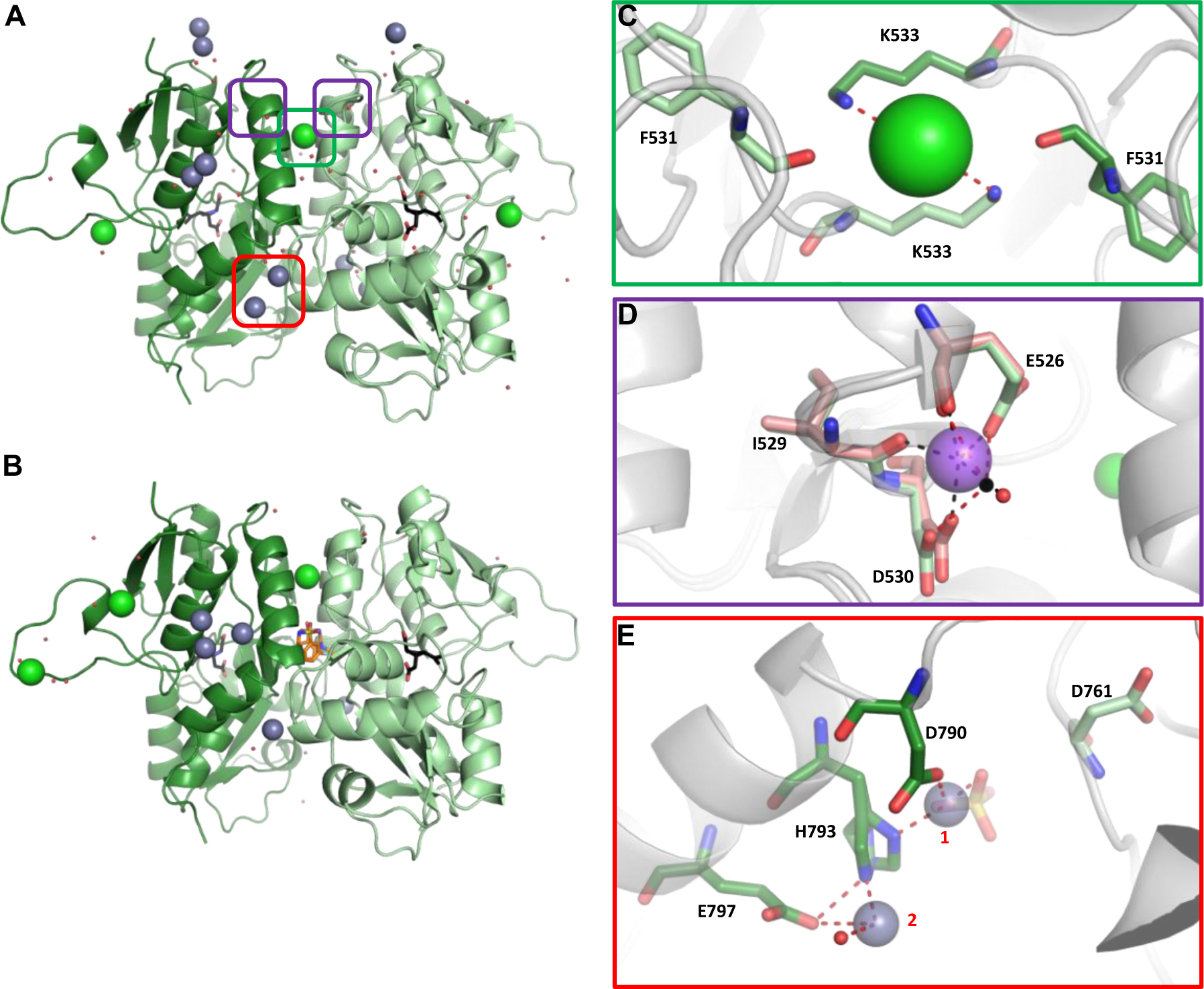
Ion binding sites at the GluK3-H523A-LBD dimer interface. (A) GluK3-H523A-LBD dimer with kainate (black carbon atoms), showing location of ions: chloride green and zinc lightblue. Chains A and B are displayed in light and dark green, respectively. Red dots represent water molecules. (B) GluK3-H523A-LBD dimer with kainate and BPAM344 (orange carbon atoms). Chains A and D are displayed in light and dark green, respectively. (C) Zoom on the chloride binding site in the GluK3-H523A-LBD dimer (grey cartoon) with kainate. Residues from the two subunits are shown in light and dark green, respectively. The chloride ion is shown as a green sphere. Potential contacts of the chloride ion are shown as red dashed lines. (D) The putative sodium ion binding site. Overlay of GluK3-H523A-LBD with kainate (green residues) on GluK1-LBD with kainate (pdb-code 3C32, chain A; salmon residues (14)). The sodium ion in GluK1-LBD is shown in violet and its contacts (black dashed lines) to the surrounding residues and a water molecule (red sphere) are indicated. In GluK3-H523A-LBD (chain D) a water molecule is shown as a black sphere and its contacts as red dashed lines. (E) Zoom on the dimer interface zinc-binding site in GluK3-H523A-LBD with kainate (chain B). Two zinc ions are shown as lightblue spheres. Potential contacts to zinc within 2.7 Å are shown as red dashed lines. See also Figure S3. Oxygen atoms are red, nitrogen atoms blue, sulfur atoms yellow, and fluorine atoms lightblue.

The GluK3-H523A-LBD dimers engage in several zinc-mediated crystal packing contacts to other GluK3-H523A-LBD dimers (Figure 7(A,B,E), Figure S3). However, one side of each dimer (chain B in the AB dimer and chain D in the CD dimer) is not engaged in any crystal packing interactions with neighboring molecules, allowing investigation of potential biologically relevant zinc binding sites (Figure 7(A,B,E), Figure S3). The electron densities indicated that these zinc ions were located close to Asp790 and His793 (Figure S3). The location of the zinc binding site is consistent with the results reported previously [13]. Zinc ion **1** coordinates to Asp790 and His793 in both chains B and D (GluK3-H523A-LBD with kainate; Figure 7(E)). Zinc ion **2** coordinating with His793 and Glu797 is observed in chain B only. In all molecules of the GluK3-H523A-LBD structures, the side chain of Asp761 of the partner subunit adopts a conformation pointing away from the zinc binding site. Therefore, we performed MD simulations to investigate if Asp761 is likely to participate in zinc binding. The simulations suggested that both Asp761 and the zinc ion coordinated by Asp790 can move to allow Asp761 to coordinate the zinc ion together with Asp790 (Figures S4 and S5).

### BPAM344 induces a compact conformation of full-length GluK3

To investigate if BPAM344 can stabilize the homomeric GluK3 in a conformation different from the structure of desensitized GluK3 with SYM (pdb-code 6JFY) [29], we purified full-length rat GluK3 with deletion of the CTD. Using electron microscopy, we found that GluK3 in the presence of glutamate alone adopts a desensitized conformation as previously observed in the presence of SYM (Figure 8). The NTD layer adopts an N-like conformation of two NTD dimers. The LBD layer has a four-fold like arrangement, and this arrangement of the LBD layer has previously been described as an inverted pyramid [26]. A dominating desensitized form of GluK3 would be consistent with fast desensitization of the GluK3 receptor in the presence of glutamate.

**Figure 8.**
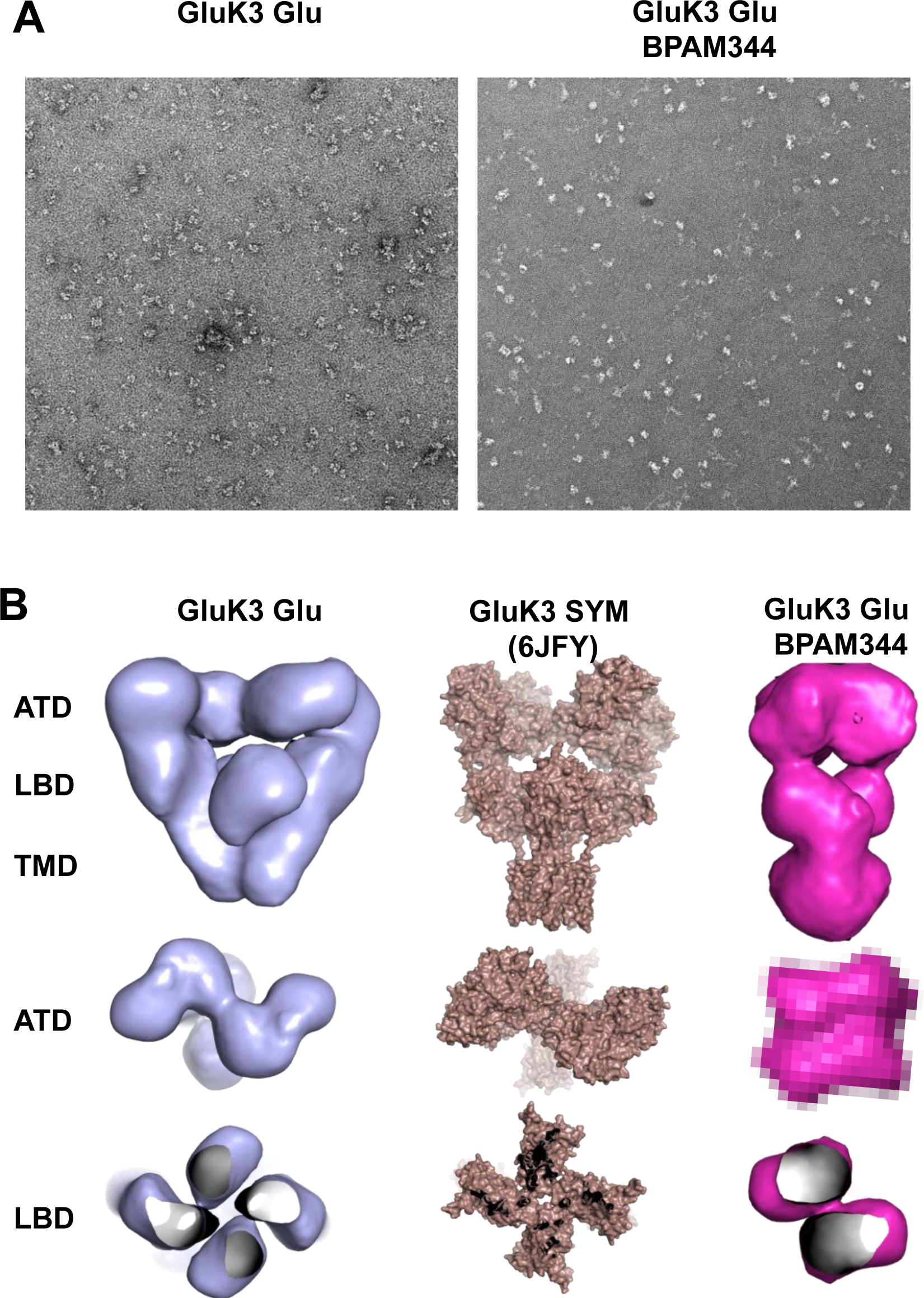
Electron microscopy of GluK3 with glutamate and glutamate/BPAM344, respectively. (A) Representative micrograph of GluK3 with glutamate (left) and glutamate/BPAM344 (right). (B) 3D reconstruction volume of GluK3 with glutamate (blue) and glutamate/BPAM344 (magenta). The structure of desensitized GluK3 with SYM (pdb-code 6JFY, brown [29]), is shown for comparison. The location of the ATD, LBD, and TMD layers are indicated in the figure.

For GluK3 in the presence of glutamate and BPAM344, we found a compact structure of GluK3 to be present in ∼30% of the conformations, whereas the desensitized form was the major conformation. The compact structure of GluK3 has a dimer-of-dimers arrangement at both the LBD and NTD layer (Figure 8). This dimer-of-dimers arrangement is similar to the the “activated” form of the AMPA receptor GluA2 [34] (Figure S6).

## Discussion

In 2017, Larsen *et al*. reported that glutamate-induced responses were significantly potentiated by BPAM344 at GluK1-3 receptors using electrophysiology [31]. This opened the opportunity to establish high-throughput calcium-sensitive fluorescence-based assays in parallel at GluK1-3. By applying BPAM344 in the assays we were able to diminish kainate receptor desensitization and thereby detect measurable agonist responses. These assays allow the determination of potencies of agonists, antagonists, and positive allosteric modulators.

Larsen *et al*. [31] reported an estimated potency of 79 µM of BPAM344 at GluK2 in electrophysiological experiments. Using the calcium-sensitive fluorescence-based assay, we found a similar EC_50_ value 75.4 µM of BPAM344 at GluK2. Our assays also allowed the determination of EC_50_ values of BPAM344 at GluK1 and GluK3, showing that BPAM344 is most potent at GluK1 (26.3 µM). Whereas BPAM344 showed the largest potentiation of responses at GluK3 receptors compared to GluK1 and GluK2 responses using electrophysiology and 100 µM of BPAM344 [31], we here show that BPAM344 has the weakest potency at GluK3 (EC_50_ of 639 µM). When compared to the potency of BPAM344 at the AMPA receptor GluA2 (EC_50_ of 0.9 µM [38]) it is evident that this positive allosteric modulator was originally designed for AMPA receptors.

Sagot *et al*. previously reported that domoate has nanomolar binding affinity at GluK1-3 [36]. No response to domoate at GluK3 in our assay was observed even at 100 µM domoate (data not shown). Others have also reported that the application of 100 µM domoate failed to elicit a response at GluK3 receptors [9]. Due to the lack of response to domoate at GluK3, domoate was instead investigated as an antagonist, and the results indicated that domoate acts as a very weak partial agonist or functional antagonist at GluK3 receptors.

Until now, only very low-resolution structures (7.4-10.6 Å) of GluK3 are available [28, 29], and it has therefore not been possible to locate any ligands in these structures. To increase the resolution and thereby enable detailed investigations of ligand and ion binding sites, we first focused on crystallizing the ligand-binding domain of GluK3 in complex with kainate in the presence of zinc ions. As positive allosteric modulators like BPAM344, as well as ions, are expected to bind at the LBD dimer interface, GluK3 must form dimers in crystals to determine such allosteric sites. The GluK3-LBD has previously been crystallized both with and without the presence of zinc ions, but in both cases, it crystallized in its monomeric form. To facilitate crystallization of the biological dimer, we mutated His523, located at the dimer interface, into an alanine to prevent this residue from interfering with crystal packing contacts through zinc ions.

In the GluK3-H523A-LBD, Ala523 is located in a hydrophobic pocket within the dimer interface. The His523 site in GluK3 has been previously investigated in functional studies, showing that mutations can have large effects on receptor function [13]. Mutation of His523 into alanine resulted in ∼2-fold slower desensitization compared to wildtype GluK3. Likewise, mutation of the corresponding Tyr521 in GluK2 [42] and Leu504 in the AMPA receptor GluA2 [43] into alanine resulted in ∼5-fold and ∼3-fold slowed desensitization, respectively. It was suggested that the Y521A mutation in GluK2 improves dimer packing, as the Tyr521-Lys525 contact is lost [42]. A tighter packing of GluK3 when an alanine is present instead of His523 is also likely. It is well-known that kainate receptors require both anions and cations for activation [44]. X-ray structures of GluK1 and GluK2 LBD dimers revealed that one chloride ion and two sodium ions bind at the top of the dimer [14, 15]. As the residues in the anion binding site of GluK1 and GluK2 are the same as in GluK3, and the GluK3-H523A-LBD was crystallized in conditions containing chloride ions, we inspected the electron density at this site. A chloride ion at this site was observed in both the structure with and without BPAM344. The residues comprising the sodium binding site are also the same in GluK1-3. A zinc-binding site has previously been identified in GluK3 based on evidence from mutational and electrophysiological data as well as structure modelling, involving Asp790 and His793 from one subunit and Asp761 from a partner subunit of the GluK3-LBD [13]. As GluK3-H523A-LBD was crystallized in the presence of zinc ions, the dimeric GluK3-H523A-LBD structures allowed direct insight into zinc binding sites and residues involved in binding. One zinc ion was observed to coordinate to Asp790 and His793 and a second to His793 and Glu797. Whereas Asp761 in the previous study [13] was found to be an important residue for zinc ion binding, we observed that Asp761 adopts a conformation pointing away from the zinc binding site in all 12 molecules of the two GluK3-H523A-LBD structures. As this could be a result of crystallization, we investigated the dynamics of Asp761 using molecular dynamics simulations. The simulations suggested that both Asp761 and the zinc ion coordinated by Asp790 can move to allow Asp761 to coordinate the zinc ion together with Asp790.

To investigate if BPAM344 can stabilize GluK3 in a compact conformation showing a dimer-of-dimers arrangement as the “activated” form of the AMPA receptor GluA2 [34] we looked at particles of purified full-length GluK3 using electron microscopy. We found that GluK3 in the presence of glutamate alone adopts a desensitized conformation, whereas GluK3 in the presence of glutamate and BPAM344 is present as a compact structure in ∼30% of the conformations, with the desensitized form as the major conformation. This compact structure of GluK3 has a dimer-of-dimers arrangement at both the LBD and NTD layer. Interestingly, whereas binding of the antagonist UBP310 led to a partially “recovered” state [29], binding of BPAM344 led to full recovery from desensitization in approximately 1/3 of the molecules. This agrees well with the observation that BPAM344 dramatically potentiates glutamate-evoked currents of GluK3 [31]. Compared to the AMPA receptor GluA2 for which a structure of an “activated” complex bound to glutamate, cyclothiazide, and stargazin in digitonin is available (pdb-code 5WEO, [34]), the dimer-of-dimers arrangement at the NTD layer seems to be more compact in GluK3 than observed for GluA2. However, higher resolution structures using cryo-EM are required to fully characterize the structural details of individual domains. Until date, only low resolution cryo-EM structures of GluK3 have been reported in the resolution range 7.4-10.6 Å [28, 29]. We have experienced similar challenges with low resolution of GluK3 as well as with solubilization of BPAM344 (high concentration required due to low potency at GluK3) for cryo-EM studies and therefore used negative stain EM requiring lower concentrations.

In conclusion, we were successful in crystallizing the ligand-binding domain of GluK3 as a dimer by introduction of a H523A mutation at the dimer interface. This allowed the determination of two positive allosteric binding sites at the dimer interface as well as potential zinc, sodium, and chloride ion binding sites. In the presence of glutamate and BPAM344, full-length GluK3 was shown to adopt a compact dimer-of-dimers arrangement using electron microscopy. The calcium-sensitive fluorescence-based assays for GluK1-3 hold the potential for a high-throughput screening tool to identify new pharmacological compounds targeting kainate receptors. This new structural insight into the GluK3 LBD dimer and full-length structure, combined with three screening assays that can be run in parallel, may contribute to unravelling the potential of kainate receptors as targets for the treatment of CNS diseases.

## Materials and Methods

### Experimental Design

In this study, we established stable expression of the kainate receptors GluK1-3 in the Grip Tite™ 293 MSR (GT-HEK293) cell line for application in calcium-sensitive fluorescence-based assays. These assays were used to characterize glutamate, kainate, domoate, UBP310, and BPAM344. For all experiments, number of samples, experiment replicates, and statistical test used are reported in the figure legends.

We also designed constructs of the ligand-binding domain of GluK3 and full-length GluK3 for use in X-ray crystallography and electron microscopy studies, respectively. Molecular dynamics simulations were applied to further investigate the cation and anion binding sites.

### Constructs for GluK1-3 stable cell lines and GluK3-LBD

The cDNAs encoding for the rat kainate receptors were amplified using the respective plasmids (GluK1(*Q*)_1b_-pCis, GluK2(*VCQ*)_a_-pGEMHE and GluK3_a_-pGEMHE) and subcloned into pIRES with either blasticidin, puromycin, or hygromycin resistance using Xbal/Xhol, BssHI/Nhel and BstEII/BssHII as restriction enzymes, respectively. GluK1(*Q*)_1b_ and GluK2(*VCQ*)_2a_ both contain a glutamine residue in their respective Q/R site, making them Ca^2+^ permeable. In addition, GluK2 contains RNA-editing at the I/V and Y/C site further enhancing the calcium-permeability of the channel [45-47].

The vector pOPINJ [48] containing the GluK3-LBD gene [16] was used as a template to generate two fragments containing the H523A mutation. Forward and reverse primers with mutated site (CCGCTGACCATTACCGCTGTGCGTGAAAAAGCG and CGCTTTTTCACGCACAGCGGTAATGGTCAGCGG) and forward and reverse fragments for a GluK3 LBD gene with appropriate extensions for cloning were used. Two fragments were further connected by PCR and the resulting insert was cloned into the pOPINJ vector using the In-Fusion cloning kit as described [48]. The presence of the mutation was confirmed by sequencing.

### Radioligand binding assays

The K_d_ of NF608 [35] and kainate was determined using homologous competition binding (Figure S1). Cells from the three generated cell lines were harvested in 0.5-1 mM EDTA/phosphate buffer saline (PBS), spun at 300 x g for 5 min, and the cell pellets were then homogenized in ice-cold 5 mM Tris-HCl buffer (pH 7.4, 4°C), centrifuged at 20,000 x g in a pre-cooled centrifuge (4°C, 10 min.). The latter step was repeated once more before the pellets were re-suspended in 50 mM Tris pH 7.1. Membranes were incubated in 5 mL glass tubes in a total volume of 250 µL with 5-20 nM [^3^H]-NF608 (16.3 Ci/mmol) or 2-20 nM [^3^H]-KA (67.5 Ci/mmol) and µM - mM unlabelled NF608 or kainate dissolved in 50 mM Tris pH 7.1. Equilibration was carried out at 4°C for 2 h and terminated by rapid filtration onto GF/C filters (pre-soaked in 0.3% polyethyleneimine). Filters were washed twice with 4 mL ice-cold 50 mM Tris pH 7.1, placed in pony vials, and 3 mL UltimaGold scintillation liquid was added. After 12 h equilibration, radioactivity was determined by scintillation counting on a PerkinElmer TriCarb 2900 scintillation counter.

### Calcium-sensitive fluorescence-based assays

Initial screening employing various types and compositions of buffers, washing volumes, and washing steps were conducted as part of optimizing the GluK1-3 assays. The stable cell lines were grown until 85-95% confluency before splitting them into poly-D-lysine-coated black/clear flat bottom 96-well plates with the construct’s respective culture medium. Subsequently, plates were incubated at 37°C in a humidified 5% CO_2_ incubator and allowed to reach near cell confluence lasting around 2-3 day. Cells were washed twice with 200 µL PBS per well with an automatic multi-channel pipette and loaded with 50 µL DMEM containing 2 µM of either Fluo-4-AM or Fluo-8-AM. Immediately after loading of the dye, cells were incubated at 37°C in humidified 5% CO_2_ incubator for at least 30 min. After incubation, the dye-solution was discarded and the cells were rinsed three times with 200 µL PBS before adding 50 µL of Fluo buffer (100 mM choline chloride, 10 mM NaCl, 5 mM KCl, 1 mM MgCl_2_, 20 mM CaCl_2_, and 10 mM HEPES pH 7.4) per well. The plates were incubated at room temperature (20-23°C) for 10 min before loading them into a FLEXstation I Plate Reader instrument. Drug solutions were prepared in Fluo buffer and 75 µL of drug solution was loaded per well into a V-bottom shaped drug plate. For conducting inhibition experiments the same steps were followed except: after the final wash the Fluo buffer contained the inhibitor alone. GluK1-3 receptor activation in the stable cell lines was measured in relative fluorescence unit (RFU) at an emission wavelength of 538 nm caused by excitation at 485 nm. Background fluorescence (zero baseline; set as 8 points) was measured for 16 sec, followed by addition of 50 µL of drug solution per well and fluorescence was measured for 104 sec, in total 2 min for the experimental time (or 3 min for GluK3). Peak fluorescence was calculated as the difference between the maximal fluorescence and the background fluorescence, i.e., resulting peak fluorescence responses were calculated by subtracting the baseline (average of 8 measurements before compound addition) for each individual curve using SoftMax Pro version 5.4 software (Molecular Devices).

### Expression and purification of GluK3-H523A-LBD

Expression and purification of the rat GluK3-H523A-LBD were essentially performed as previously described [16]. Briefly, the protein was expressed in *E. coli* and after cell disruption the cleared lysate was applied to a GSTrap chromatography column (GE Healthcare). After the on-column cleavage with 3C protease (prepared according to in house protocol), the GluK3-H523A-LBD was eluted and further concentrated and separated by size exclusion chromatography using a Superdex 75 column (GE Healthcare).

### Crystallization of GluK3-H523A-LBD

Crystals were obtained by the hanging drop vapor diffusion method at room temperature. The solution of GluK3-H523A-LBD with kainate consisted of ∼4 mg/mL of GluK3-H523A-LBD and 17 mM kainate (in 10 mM HEPES pH 7.0, 150 mM NaCl, 1 mM EDTA, and 10% glycerol). For crystallization of GluK-H523A-LBD with kainate and BPAM344, the protein– ligand solution consisted of ∼4 mg/mL of GluK3-H523A-LBD, 6 mM kainate, and a suspension of BPAM344. The suspension of BPAM344 was prepared by soaking 1.6 mg BPAM344 in 1.3 mL water and 2 µL 1 M NaOH.

The crystallization drops were made by mixing equal volumes (1 µL) of protein and reservoir solution (15% PEG4000, 0.3 M lithium sulfate, 0.1 M Tris pH 8.5, and 5 mM zinc acetate). Streak seeding was used to achieve crystals suitable for data collection. Crystals were flash cooled in liquid nitrogen using a cryo-protecting solution (20% glycerol, 20% PEG4000, 0.3 M lithium sulfate, Tris pH 8.5, and 5 mM zinc acetate).

### X-ray structure determination of GluK3-H523A-LBD

X-ray diffraction data were collected at the MAX-lab beamline I911-3, Lund, Sweden [49]. The data were processed in XDS [50], scaled, and merged in AIMLESS [51] or Scala [52], and the two structures determined by molecular replacement in Phaser [53] within the CCP4 programme suite [54]. The structure of GluK3-LBD in complex with kainate (pdb-code 4E0W, chain A [17]) was used as search model. The structure models were rebuilt using AutoBuild [55] within Phenix [56]. The structures were manually rebuilt in Coot [57] and refined in Phenix. Atomic coordinates for BPAM344 were created in Maestro (Maestro version 2.3; Schrödinger, LLC: New York, 2013) and geometry optimized (MacroModel 10.1; Schrödinger, LLC: New York, 2013). A restraint file for BPAM344 was generated using eLBOW [58], keeping the geometry from Maestro. Non-crystallographic symmetry (NCS) and TLS (1 group per chain) were applied during refinements. B-factors were refined isotropically (B-individually for GluK3-KA, B-group for GluK3-KA-BPAM344). Figures were prepared in PyMol (The PyMOL Molecular Graphics System, Version 2.0.3, Schrödinger, LLC).

### Expression and purification of GluK3

The rat GluK3 construct was subcloned from the pGEMHE vector. The GluK3 construct was modified to exclude the C-terminal part (end with EFIYKLRK) and to include a C-terminal GFP-His tag and 3C protease cleavage site. The construct corresponds to wildtype GluK3 except for a P271H mutation in the NTD. After cleavage, additional nine residues were left from the tag (SSGLEVLFQ). The GluK3 gene as well as DNA coding for 3C-GFP-His tag were amplified separately and then connected by PCR using both fragments. The resulting product was cloned into a pBAC-6 vector (Merck) by In-Fusion cloning (Clontech). The resulting vector was further used for transfection with BD BaculoGold (BD Biosciences) and virus generation in *Sf*9 insect cells. The GluK3 construct was checked by sequencing (Eurofins Genomics).

The GluK3 gene cloned into pBAC-6 was expressed in *Sf*9 insect cells using baculo virus. For purification, cells were harvested by centrifugation and resuspended in lysis buffer (20 mM HEPES pH 8.0, 300 mM NaCl, and 50 mM glutamate). Cells were lysed using sonication and membranes were harvested by centrifugation at 41,800 rpm for 1 hour. Membranes were solubilized by douncing in solubilization buffer (20 mM HEPES pH 8.0, 300 mM NaCl, 2% DDM detergent (Anatrace D310S), 50 mM glutamate, and protease inhibitor cocktail (Roche)) and stirred for 2 hours. Insoluble material was removed by centrifugation and the supernatant containing the receptor was rotated with Talon resin (Takara) overnight. Resin was loaded onto a column and washed with wash buffer (20 mM HEPES pH 8.0, 300 mM NaCl, 0.05 % DDM (Anatrace D310), 50 mM glutamate, protease inhibitor cocktail, and 25 mM imidazole). Protein was eluted in the same buffer adjusted to 250 mM imidazole. The His-tag was removed by adding 3C protease and the digested receptor was isolated using size exclusion chromatography (SEC) on a Superose 6 column in SEC buffer (20 mM HEPES pH 8.0, 150 mM NaCl, 0.05 % DDM detergent (Anatrace D310), and 50 mM glutamate).

### Electron microscopy

The diluted GluK3 sample (0.1 µM GluK3, 25 mM glutamate, and 0.1 mM BPAM344 in 10 mM HEPES pH 8.0, 150 mM NaCl, and 0.05% DDM sol. grade (0.5 mg/ml)) was applied on a 400 mesh glow discharged collodion and carbon coated copper grid and stained with 2% uranyl formate. A total of 537 micrographs were collected at 120 kV using a Tecnai G2 Spirit TWIN electron microscope with a defocus value of 0.7-1.7 µm. Images were automatically collected using a Tietz TemCam-F416 CMOS camera at a nominal magnification of 67,000x and a pixel size of 1.57 Å (binned to 3.14 Å), employing Leginon [59]. The micrographs were screened using the program Xmipp [60]. A total of 456 micrographs were selected and phase flipped for further analysis in CryoSPARC [61]. A total of 52,178 particles were extracted (box size 100x100 pixels) after blob picking with diameters between 200 Å and 300 Å and input to 2D classification. Three successive runs of 2D classification were undertaken with 50 classes. In each run particles belonging to blurry classes with low population, or clearly representing other molecules such as GROEL, were removed from the dataset leaving a total of 28,717 particles. Five *ab initio* volumes were calculated in symmetry *C*1 and all used as input for heterologous refinement with *C*2 symmetry imposed. After non-uniform refinement of the volumes, they were compared visually in ChimeraX [62]. Particles from one of the volumes were discarded as it had clearly collected bad particles. Another round of heterologous and non-uniform refinement was undertaken again using five volumes. The resulting volumes could be divided in two groups: one with a quadratic arrangement of the LBD layer characteristic of a desensitized receptor, derived from 15,528 particles and one with an arrangement of the LBDs as dimer-of-dimers, derived from 7,253 particles. Resolutions of 6.7 Å were calculated using the FSC gold standard for both volumes.

For GluK3 with glutamate only, the data were collected and processed using a protocol similar to that used for GluK3 with glutamate and BPAM344. A total of 546 micrographs were selected and phase flipped for further analysis in CryoSPARC. A total of 82,515 particles were extracted (box size 100x100 pixels) after blob picking with diameters between 200 Å and 300 Å and input to 2D classification. Four successive runs of 2D classification were undertaken with 50 classes in the first three and 20 in the last. In each run particles belonging to blurry classes with low population, or clearly representing other molecules such as GROEL, were removed from the dataset leaving a total of 19,116 particles. Three *ab initio* volumes were calculated in symmetry *C*1 and all used as input for heterologous refinement with *C*2 symmetry imposed. After non-uniform refinement of the volumes, they were compared visually in ChimeraX. The resulting volumes all had a quadratic arrangement of the LBD layer characteristic of a desensitized receptor. Therefore, all particles were used in a final round of non-uniform refinement with the lowest resolution volume from the heterologous refinement as starting volume. A resolution of 6.6 Å was calculated using the FSC gold standard.

### Molecular dynamics simulations

For MD simulations, a close to fully refined dimer of the GluK3-H532A-LBD was used, containing kainate in the two agonist binding sites, either two or four zinc ions, one chloride ion as well as two potential sodium ions that were modelled according to the position in the structure of GluK1 (pdb-code 3C32 [14]). Water molecules and other ions not located at the dimer interface were deleted. During simulations zinc, chloride, and sodium ions were retained or deleted according to the different setups indicated below and in Figure S2.

For simulations of the wildtype GluK3 LBD dimer, Ala523 in the crystal structure was mutated to histidine using the mutate function of PyMOL (The PyMOL Molecular Graphics System, Version 1.4, Schrödinger, LLC). Simulated complexes include wildtype GluK3-LBD and GluK3-H523A-LBD with no bound ions, with the two zinc ions bound to one subunit and with two zinc ions bound to both subunits; always with kainate bound at the agonist site. In addition, GluK3-H523A-LBD was simulated with the upper or lower zinc ions, respectively, bound in both chains. The complexes were inserted into a water box of dimensions (110 Å)^3^, and the systems were neutralized and 150 mM NaCl was added. MD simulations were performed using Gromacs 5.0.2 [63] with the AMBER99SB-ILDN force field [64] used for the protein along with the TIP3P water model [65] for water molecules. Parameters for kainate were generated with the general AMBER force field (GAFF v 1.7) [66]. Hydrogen atoms for kainate were added in Maestro v 9.7 (Schrödinger, New York, NY; academic version) after which parameters were generated using the antechamber module of AmberTools14 [67], employing AM1-BCC charges [68]. The AMBER format was converted to the GROMACS format using ACPYPE [69].

The simulated systems were energy minimized for 50,000 steps or until the maximum force on an atom was less than 100 kJ/mol/nm. The minimization was followed by a 200 ps equilibration in the NVT ensemble at a temperature of 300 K, controlled by a Berendsen thermostat [70]. Periodic boundary conditions were employed and van der Waals interactions were cut off at 10 Å. The long-range electrostatics were accounted for by the Particle-Mesh Ewald method [71] and all bonds were treated as constraints using the LINCS algorithm [72], allowing a time step of 2 fs. Subsequently, 1 ns of equilibration in the NPT ensemble was performed, at a pressure of 1 bar maintained by a Berendsen barostat [70]. For both steps of the equilibration, the protein heavy-atoms were position restrained with a force constant of 1000 kJ mol^-1^ nm^-2^. Finally, 100 ns of production run was performed in the NPT ensemble with settings as above but with all position restraints removed. Two repeats were run for each setup. The trajectories were analyzed using analysis tools of Gromacs and VMD [73].

### Statistical Analysis

For radioligand binding, all experiments were performed in triplicate and data were analyzed using GraphPad Prism (version 9.3.0, San Diego, CA) utilizing one site-homologous equation for [^3^H]-KA and [^3^H]-NF608. K_d_ values are represented as mean ± SEM.

In the calcium-sensitive fluorescence-based assays, all experiments were performed in octuplicates and are shown as mean ± SEM from at least three individual experiments. For each concentration-response curve, the response has been normalized to the response at maximal compound concentration. GraphPad Prism (version 9.3.0, San Diego, CA) was used for curve fitting by applying the equation: log(agonist) vs. response – variable slope (four parameters) for determining the EC_50_ and the equation: log(inhibitor) vs. response – variable slope (four parameters) for determining the IC_50_ with top constrained to 100% and bottom to 0%. Data representations in histograms and expressed with statistical analysis were performed with SigmaPlot (version 14.0, Systat Software, San Jose, CA).

## CRediT authorship contribution statement

**Yasmin Bay, Maria Musgaard**: conceptualization, methodology, investigation, formal analysis, validation, visualization, writing - original draft. **Raminta Venskutonytė, Stine M. Frantsen, Thor S. Thorsen**: conceptualization, methodology, investigation, formal analysis, validation, writing - original draft. **Karla Frydenvang**: investigation, formal analysis, validation, visualization, writing - review & editing. **Pierre Francotte, Bernard Pirotte**: resources, writing - review & editing. **Philip C. Biggin**: conceptualization, methodology, formal, analysis, validation, supervision, funding acquisition, writing - review & editing. **Anders S. Kristensen**: methodology, formal analysis, validation, visualization, supervision, writing - review & editing. **Thomas Boesen**: methodology, validation, supervision, writing - review & editing. **Darryl S. Pickering**: conceptualization, methodology, investigation, formal analysis, supervision, writing - review & editing. **Michael Gajhede**: conceptualization, methodology, investigation, formal analysis, validation, writing - original draft. **Jette S. Kastrup**: conceptualization, methodology, investigation, formal analysis, validation, visualization, supervision, funding acquisition, writing - original draft.

## Data availability

The structure coordinates and corresponding structure factor files of the GluK3-H523A-LBD dimer have been deposited in the Protein Data Bank (PDB) under the accession codes 8BST (kainate) and 8BSU (kainate and BPAM344). The maps of full-length GluK3 have been deposited in the Electron Microscopy Data Bank (EMDB) with entry ID EMD-15985 (glutamate) and entry ID EMD-15986 (glutamate and BPAM344).

## Abbreviations

AMPA: α-amino-3-hydroxy-5-methylisoxazole-4-propionate
ATD: amino-terminal domain
con A: concanavalin A
DA: domoate
Glu: glutamate
GluK1-LBD: ligand-binding domain of GluK1
GluK3-LBD: ligand-binding domain of GluK3
GluK3-H523A-LBD: ligand-binding domain of GluK3 with introduction of H523A mutation
iGluRs: ionotropic glutamate receptors
KA: kainate
LBD: ligand-binding domain
MD: molecular dynamics
RFU: relative fluorescence unit

## Acknowledgments

Heidi Nielsen and Heidi Peterson are thanked for technical support. We acknowledge the MAX lab (now MAX IV Laboratory) for time on Beamline 911-3 under Proposal MX20130012 and thanks beamline scientists for their support. Research conducted at MAX IV, a Swedish national user facility, is supported by the Swedish Research council under contract 2018-07152, the Swedish Governmental Agency for Innovation Systems under contract 2018-04969, and Formas under contract 2019-02496”. We acknowledge the European Synchrotron Radiation Facility for provision of synchrotron radiation facilities, and we would like to thank beamline scientists for assistance in using beamline ID23eh2 (project no. mx1691). We acknowledge Diamond Light Source for time on Beamline I03 under Proposal mx11011-1 and the support from beamline scientists. We thank the ARCHER U.K. National Supercomputing Services for computer time granted via the High-End Computing Consortium for Biomolecular Simulation, HECBioSim (http://www.hecbiosim.ac.uk), supported by EPSRC (EP/ 029407/1) (M.M., PCB). Research funding is acknowledged from The Lundbeck foundation (RV, SMF, JSK), The Independent Research Fund Denmark – Medical Sciences (YB, RV, TST, JSK), Danscatt (RV, TST, MG, JSK), BioStructX (RV, TST, MG, JSK), and The Alfred Benzon Foundation (MM).

## Appendix A. Supplementary data

Supplementary data to this article can be found online at (DOI to be inserted).

