## Supplementary information for "Small molecule positive allosteric modulation of homomeric kainate receptors GluK1-3: Development of screening assays and insight into GluK3 structure"

**This PDF file includes:**

Figs. S1 to S6

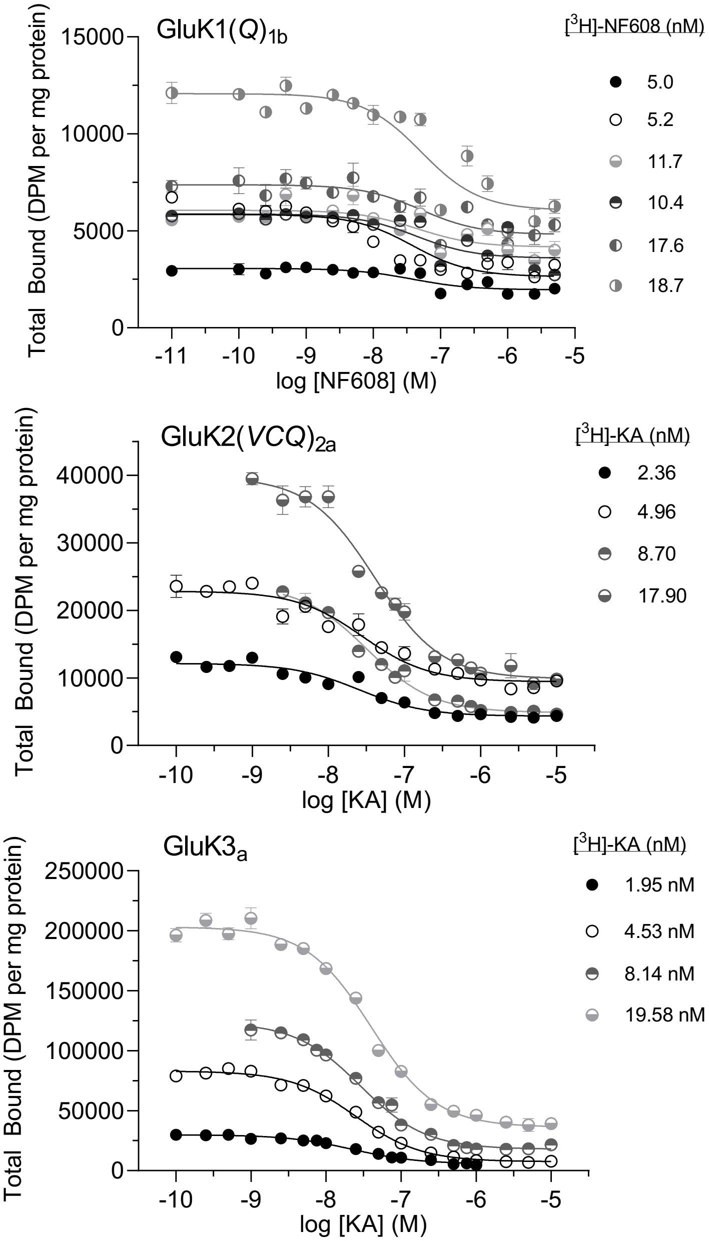


**Fig. S1. Homologous competition binding with [$^{3}$H]-NF608 as radiolabel for the GluK1(*Q*)$_{1b}$ cell line and [$^{3}$H]-kainate as radiolabel for GluK2(*VCQ*)$_{2a}$ and GluK3$_{a}$.** GluK1-3 were expressed in GT-HEK293 cells. Pooled data shown as mean ± SEM from independent experiments conducted in triplicate. In the homologous binding experiment, ligand affinity ($K_d$) is determined by analyzing the competition of varying concentrations of unlabeled ligand for four to six concentrations of labeled ligand [van Zoelen, E. J. (1992). Analysis of receptor binding displacement curves by a nonhomologous ligand, on the basis of an equivalent competition principle. *Anal. Biochem*. **200**, 393-399]. The $K_d$ values for kainate at GluK2 and GluK3 were 23 nM [CI 19-29] and 20 nM [CI 18-23], respectively. The $K_d$ value for NF608 at GluK1 was 33 nM [CI 21-53].

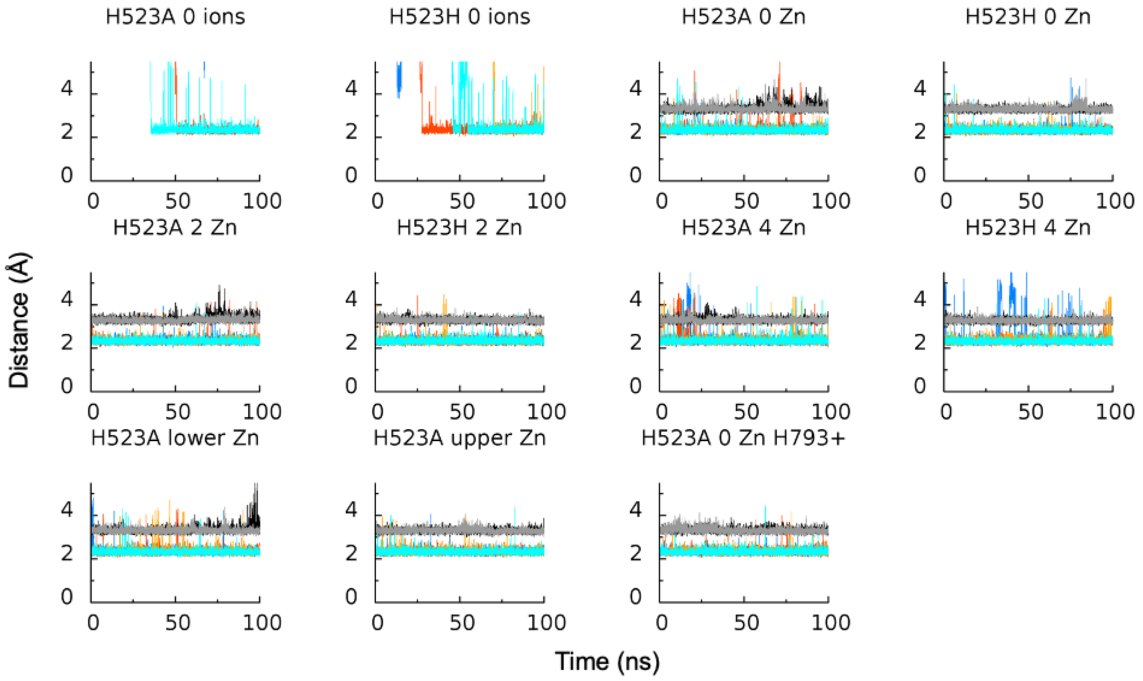


**Fig. S2. MD simulations indicate conservation of sodium and chloride sites in GluK3-LBD.** The plots illustrate distances from binding site residues to chloride and sodium ions under different simulation setups, using either the H523A as in the crystal structure or the wild-type sequence (A523 mutated back to His; denoted H523H). "0 ions" indicate that no ions were bound to the protein at the start of the simulation. "0 Zn", "2 Zn", "4 Zn", "upper Zn", and "lower Zn" refer to the zinc ions present at the start of the simulation. "H793+" indicates that H793 was treated as positively charge (if not mentioned, it is treated as neutral). Color coding: orange and blue: minimum distance (Å) from each E526 to the nearest sodium ion. Black: minimum distance from either K533 to the nearest chloride ion. Darker colors for one repeat, lighter for the second repeat. Ions appear to remain stably bound in all simulations, and some ions even find the binding sites when not present initially.

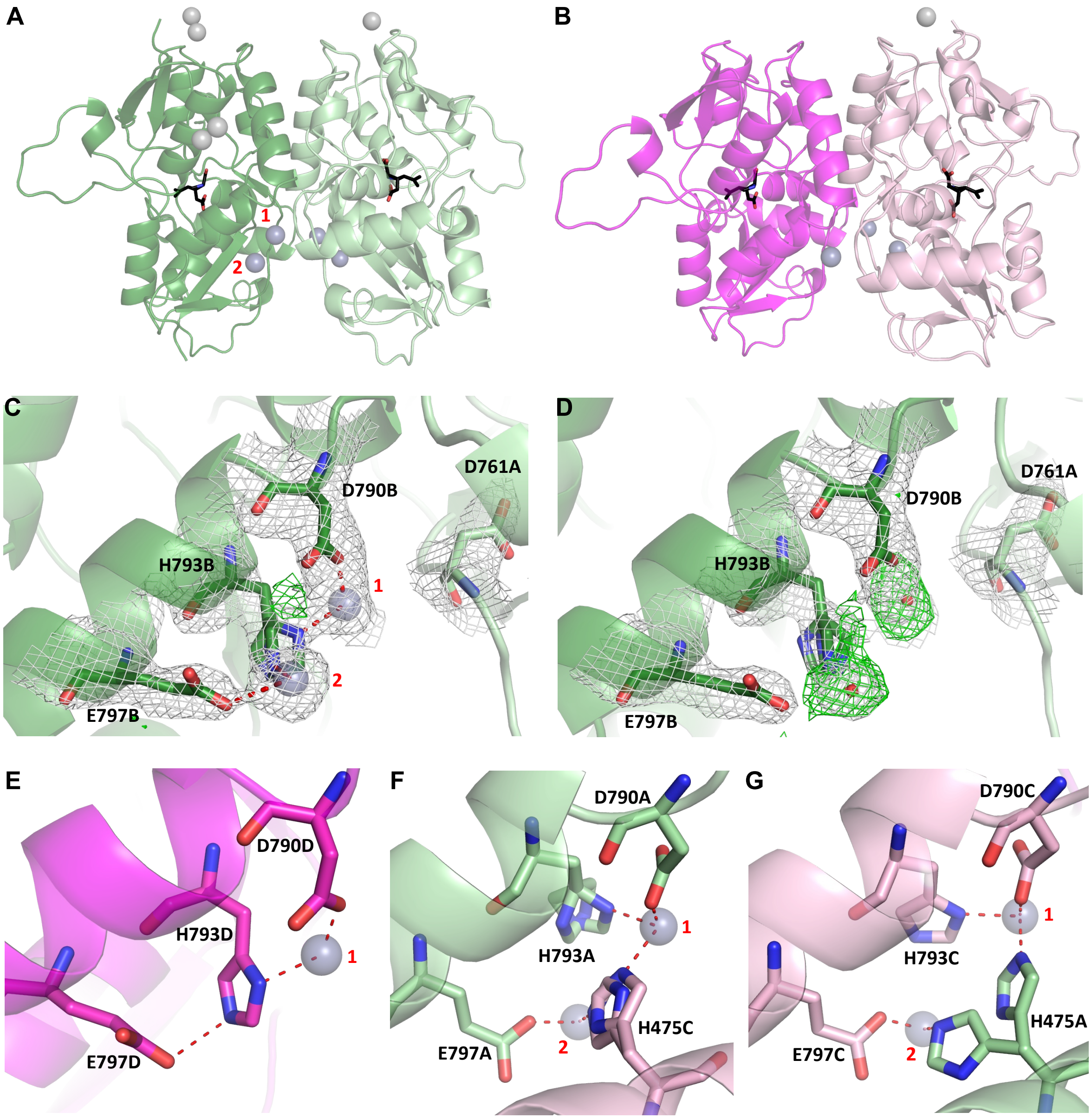


**Fig. S3. Zinc binding sites in GluK3-H523A-LBD with kainate.** (**A**) GluK3-H523A-LBD dimers with kainate (black carbon atoms), showing location of zinc ions (at dimer interface: light blue; at surface: light grey). AB dimer with chain A in light green and chain B (left) in dark green. (**B**) CD dimer with chain C in light magenta and chain D in dark magenta. (**C**) Zoom on zinc sites 1 and 2 at the dimer interface (chain B). The final 2Fo-Fc (1 sigma) and Fo-Fc (3 sigma) electron densities for the zinc atoms and surrounding residues (in stick representation) are shown. Nitrogen atoms are blue and oxygen atoms red. Potential contacts to zinc within 2.7 Å are shown as red dashed lines. (**D**) 2Fo-Fc and Fo-Fc electron densities when refining the structure with a water molecule at the zinc sites, indicating additional electrons. (**E**) Zoom on zinc site 1 at the dimer interface (chain D). (**F**) Zoom on zinc sites 1 and 2 at the dimer interface (chain A), involving crystal packing contacts with chain C. (**G**) Zoom on zinc sites 1 and 2 at the dimer interface (chain C), involving crystal packing contacts with chain A.

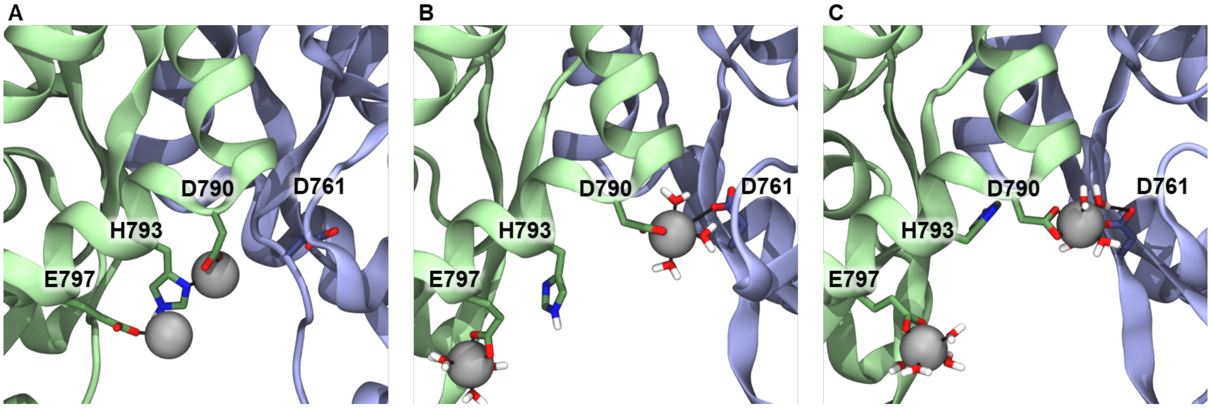


**Fig. S4. Potential coordination of zinc ions using MD simulations**. (**A**) Conformation from the crystal structure. (**B**) Snapshot after around 13 ns of MD simulation of GluK3-LBD-H523A with kainate (second repeat) with two zinc ions bound. Asp761 seems to coordinate the upper zinc ion directly. (**C**) Snapshot after around 35 ns for the same MD simulation as in (B), where the interactions between Asp761 and the upper zinc ion are now water mediated.

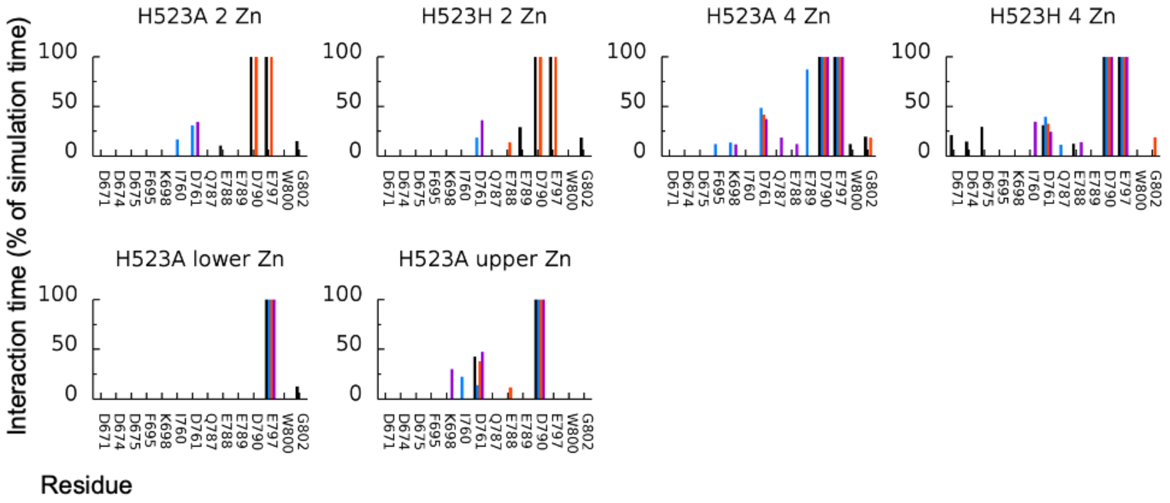


**Fig. S5. Residues involved in interactions with zinc ions during MD simulations on GluK3-LBD using either the H523A as in the crystal structure or the wild-type sequence (A523 mutated back to His; denoted H523H).** “2 Zn”, “4 Zn “, upper Zn”, and “lower Zn” refer to the zinc ions present at the start of the simulation. An interaction is defined as a distance <4 Å between the residue (non-hydrogen atoms) and a zinc ion. The interaction time is given as percentage of time over the trajectory. Black and blue represent chain A and chain B, respectively, of the first repeat while red and purple represent chains A and B, respectively, of the second repeat.

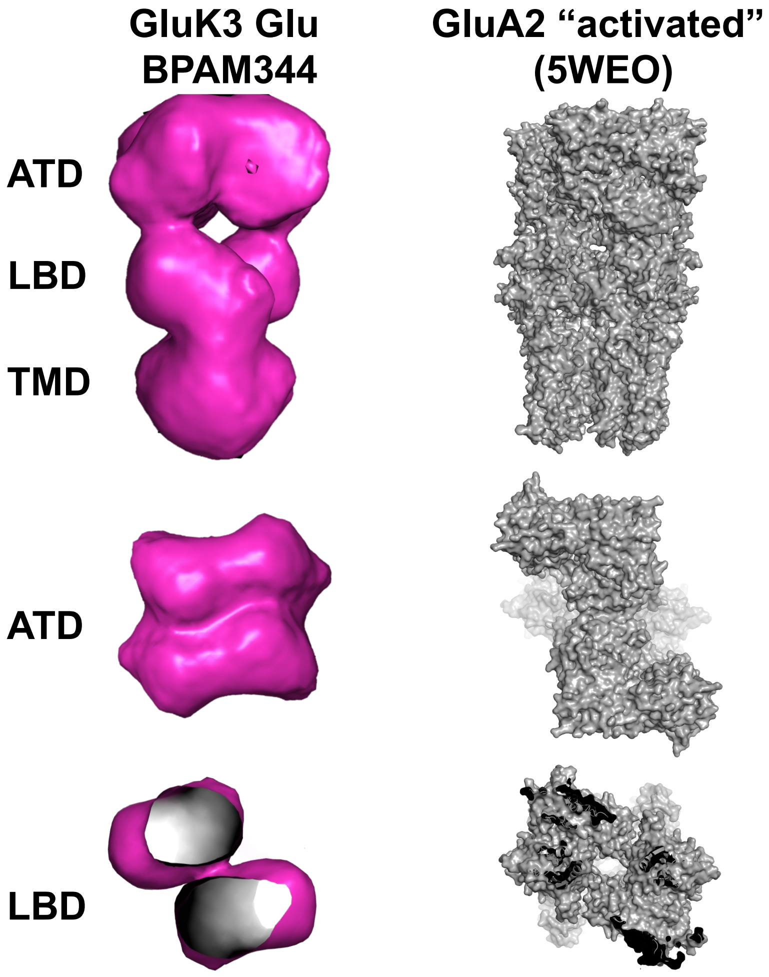


**Fig. S6. Electron microscopy of GluK3 with glutamate/BPAM344**. Left: 3D reconstruction volume of GluK3 with glutamate/BPAM344. Right: The structure of the “activated” GluA2 bound to glutamate, cyclothiazide, and stargazin in digitonin (pdb-code 5WEO) is shown for comparison. The location of the ATD, LBD, and TMD layers are indicated in the figure.
